# Neonatal personality is associated with sex-specific developmental trajectories in a wild mammal

**DOI:** 10.64898/2026.11.06.764900

**Authors:** Cong Yu, Simone Ciuti

## Abstract

Animal personality, defined as consistent among-individual differences in behaviour, is an important source of phenotypic variation, but its developmental origins and associations with subsequent development remain poorly understood. We measured neonatal personalities, early-life growth, and yearling male antler length in wild fallow deer (*Dama dama*). Using eight years of data, we estimated repeatability and among-individual correlations among neonatal personalities and growth and applied structural equation modelling to examine associations with yearling male morphology. Heart rate and behavioural reaction measures showed repeatable among-individual variation, demonstrating that stable differences were detectable shortly after birth. Behavioural reaction was negatively correlated with growth in males but not in females. Heart rate was not associated with growth in either sex. In males, the effects of neonatal behavioural reactivity persisted beyond early life: more behaviourally reactive neonates had shorter antlers as yearlings, indirectly through reduced early-life growth. These findings show that stable among-individual differences emerge during the neonatal period and are associated with sex-specific developmental trajectories.

## Introduction

Consistent individual differences in behaviour, or animal personalities, are a fundamental source of intraspecific variation with profound ecological and evolutionary consequences (Sih et al., 2004; Wolf & Weissing, 2012). This variation is often structured into behavioural syndromes, in which suites of behaviours are correlated across contexts (Sih et al., 2004). Such syndromes are increasingly recognised as key drivers of individual fitness, influencing survival, resource acquisition, and reproductive success (Briffa, Sneddon, & Wilson, 2015; Réale et al., 2009; Smith & Blumstein, 2008). Consistent with Réale, Dall and Dingemanse (2026), examining these components of behavioural variation can reveal how traits are structured and how phenotypes are organised during development.

However, a critical and largely unresolved question concerns their developmental origins and whether these early differences are associated with subsequent growth and morphological development. This question is particularly difficult to resolve in large, long-lived herbivores, where the link between early-life condition and later phenotype is often obscured. Individuals can show large differences in body mass among cohorts at birth, but this variation may dissipate with age as individuals exhibit compensatory “catch-up” growth (Hamel et al., 2016). This raises a critical question: are behavioural differences expressed during the neonatal period associated only with early development, or are they also linked to later morphology? Understanding the link between early-life personality and growth has broad implications for ungulate ecology and evolution. It may reveal how early-life behavioural variation is associated with developmental trajectories, population dynamics, and individual differences in fitness.

These developmental associations may differ between females and males. The sexes often follow distinct growth trajectories and may therefore differ in how neonatal behavioural and physiological variation is associated with subsequent growth and morphology. Empirical studies support this possibility where the strength and direction of covariation among behaviour, physiology, body condition, and structural size can vary between populations and sexes (Gaona-Gordillo et al., 2023). This question is especially relevant in sexually dimorphic ungulates because males face stronger selection to grow rapidly and attain a large body size, so early phenotypic differences may be more strongly associated with their subsequent growth (Post et al., 1999). Examining females and males separately therefore allows us to determine whether behavioural differences expressed at birth are associated with sex-specific patterns of growth and whether these associations are stronger in males.

Here, we investigate the link between neonatal personality and early-life development in a large, sexually dimorphic mammal, the fallow deer (*Dama dama*). As a hider species, fallow deer neonates are typically hidden in tall grasses or shrubs for the first weeks of their lives and experience a period of relative isolation. Because fallow deer neonates experience limited conspecific interactions during their hider phase (Chapman & Chapman, 1997; Lent, 1974), this system allows behavioural differences to be measured before extensive social experience accumulates and their associations with subsequent developmental trajectories to be examined.

We specifically test (1) whether physiological (heart rate) and behavioural responses recorded during the neonatal period contain repeatable among-individual variation and are correlated at the among-individual level; (2) whether these neonatal responses are associated with early-life growth in females and males; and (3) in males, whether neonatal responses and early-life growth are associated with antler length at one year within a hypothesised structural model. We expect greater behavioural and physiological reactivity to be associated with lower early-life growth, with stronger associations in males than in females. We further expect greater neonatal reactivity and slower early-life growth to be associated with shorter yearling antlers. Together, these analyses test whether stable among-individual differences in neonatal behavioural and physiological responses are associated with sex-specific developmental trajectories and later male morphology.

## Methods

### 1. Study site and study species

This study was conducted in Phoenix Park, a 7.07 km² urban park in Dublin, Ireland (53.3559° N, 6.3298° W). The vegetation of the park consisted of 80% open grassland and 20% mixed woodland. During late summer, approximately 600 free-ranging fallow deer inhabit the park following the fawning period, with most neonates born in June each year (Griffin et al., 2022). As a hider species, fallow deer neonates are typically hidden in tall grasses or shrubs, separate from the main doe herd during their first weeks of life, and they are later reintegrated into the doe herd (Amin, Jennings, et al., 2022; Chapman & Chapman, 1997; Ciuti et al., 2006). The red fox (*Vulpes vulpes*) is the park’s only natural predator, but neonate predation by unleashed pet dogs (*Canis lupus familiaris*) brought by visitors has also been recorded (Conteddu et al., 2023).

## 2. Data collection

### 2.1. Neonatal personality and growth rate

As part of the ongoing monitoring and management operations of the deer population, our research group captured and tagged of neonates daily during June from 2018 to 2025 with routine patrols of traditional fawning sites utilised by does (Griffin et al., 2022). Approximately 15 trained members, supervised by an experienced wildlife biologist, used circular fishing nets (1–1.5 m in diameter; various brands) to capture the neonates (Amin et al., 2021). When encountering an untagged neonate, the team captured, ear-tagged, and recorded its neonatal response measures and age, subsequently measuring, weighing, and releasing it. The capture protocol and procedures have been described in detail in supplementary methods – but also see (Amin, Jennings, et al., 2022; Amin et al., 2021; Yu et al., 2025). We avoided recapturing fawns on the same day (mean interval between captures and recaptures was: 4.56 ± 3.40 days). We evaluated neonatal personality using a physiological metric—pre-release heart rate, reflecting the neonate’s physiological response to human handling—and a behavioural metric—the reactions upon release, indicating the neonate’s behavioural response to handling. We measured heart rate immediately prior to weighing by counting the number of beats over a 20-second period using a Lightweight Dual Head Stethoscope (MDF®, California, USA). After weighing the neonate, we opened the bag to release it and immediately recorded its initial reaction using a camera (Canon Compact PowerShot SX500 IS) for 30 seconds (i.e., the first reaction to human handling). To minimise disturbance, all team members remained still and silent once the bag was open. We categorised the neonate’s first reaction on a five-point ordinal scale: (1) remained in the bag for 30 seconds, (2) stood up but did not move, (3) took a short walk and stopped, (4) hopped out of the bag and stopped within two meters, (5) ran after release (Fig. 1). We originally distinguished between neonates that ran but stopped within sight and those that ran out of sight without stopping. However, as we did not have a standardised measure of long-distance visibility at the release site, determining whether a running neonate had truly disappeared or merely moved beyond an arbitrary visual threshold was too subjective. We therefore merged these two categories into a single level, yielding the final five-point scale. If a neonate remained in the bag for the full 30 seconds, we prompted it to leave by making a sound or gently touching it and recorded its subsequent behaviour as the second reaction. To integrate both prompted and spontaneous behavioural reaction into a single measure, second reaction was scored on a combined 9-point ordinal scale (1-9). Neonates that required prompting were scored 1-5 based on their response to the prompt (1 = remained stationary despite prompting; 5 = ran after prompting), while neonates that exited spontaneously within 30 seconds were assigned scores 6-9 based on their first behavioural reaction (6 = stood up spontaneously; 9 = ran after release), reflecting that spontaneous departure inherently represents a more reactive response than a prompted one. Previous studies quantified neonatal behaviour using latency to leave, a metric correlated with trappability (Yu et al., 2025) and resource acquisition (Amin, Jennings, et al., 2022). In this study, however, we used a five-point ordinal reaction score to characterise the neonate’s response upon release, because our aim was to capture variation in the type and intensity of the behavioural response rather than response timing alone.

**Figure. 1.**
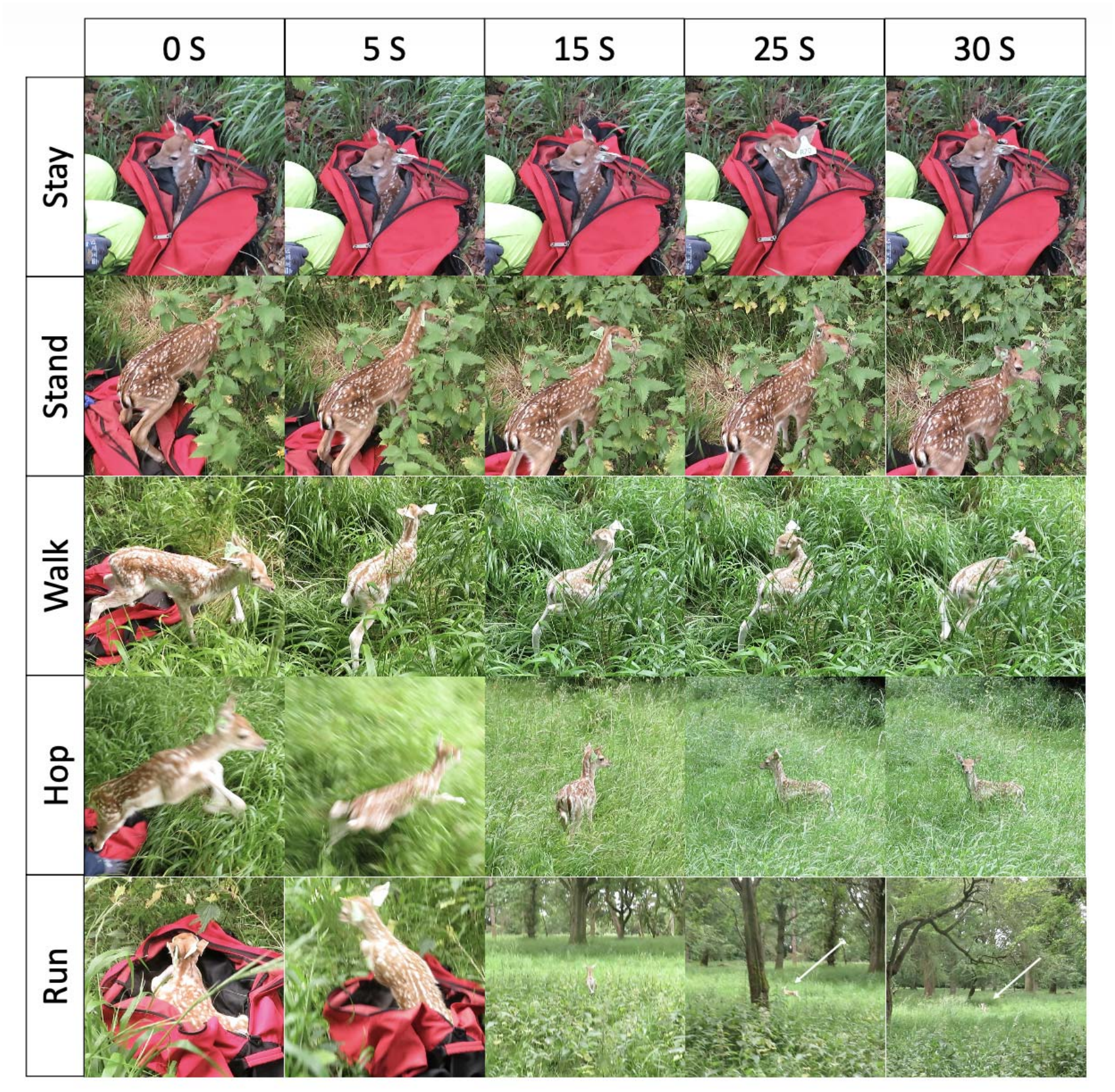
The five-point ordinal scale used to assess neonatal behavioural reactions to human handling in fallow deer neonates. We reported here the video frame captures (0, 5, 15, 25 and 30 secs, respectively) from videos taken on five neonates displaying varying behaviours at release. A score of 1 indicates that the neonate remained in the bag for 30 seconds. A score of 2 indicates that the neonate stood up without any movement. A score of 3 reflects the neonate taking a short walk before stopping. A score of 4 is assigned when the neonate hopped out of the bag and stopped within two meters, and a score of 5 is given when the neonate ran.

We recorded covariates known to influence neonatal responses to handling, including capture time (hours), handling duration (seconds), neonate sex, and the number of capture events (i.e., how many times a given fawn was captured during the fawning season) (Amin et al., 2021; Yu et al., 2025). We measured air temperature at the bed-site using a digital thermometer (Grandbeing, China). Before capture, we assessed each neonate’s prior behaviour (vigilance) upon the team’s arrival, scoring it on a scale from 0 (inactive) to 1 (active), following Amin et al. (2021). To quantify bed-site visibility, we placed a cardboard model (45 cm tall, 36 cm wide) resembling a standing neonate at the discovery location. From a distance of 10 m in each of the four cardinal directions, we recorded the percentage of unobstructed triangular areas visible on the cardboard. Observations were taken from an eye height of 70 cm, approximating the perspective of a red fox, the neonate’s primary predator at the study site (Bongi et al., 2008).

To assess growth rate, we weighed each neonate (kg) using a digital scale and a 100 L bag (resolution: 0.01 kg, Dario Markenartikelvertrieb, Hamburg, Germany). We estimated age (days) based on umbilical cord condition and hoof and dewclaw development (Galli et al., 2008). If a neonate was resighted at least 48 hours later, we recaptured and remeasured its weight and age. Each neonate was captured up to four times to minimise disturbance.

### 2.2 Proxy for body condition (antler length)

We assessed male body condition at age one by measuring investment in secondary sexual traits, i.e., antler length, a reliable indicator of phenotypic quality in fallow deer (Ciuti & Apollonio, 2011). Using photogrammetry, a validated non-invasive method (Smith, Bongi, & Ciuti, 2021), we measured antler length from the highest point to the lowest point of the burr, where it connects to the skull (Ciuti & Apollonio, 2011). Note that at this age, males have a one-spike antler with no ramifications. We collected photogrammetry data on males each August from 2019 to 2025, photographing individuals repeatedly over multiple days within the same season and across years (see (Smith, Bongi, & Ciuti, 2021) for full details on this calibrated and validated non-invasive technique). We used average value of left and right antler lengths, and, because in many instances we recorded the same antlers multiple times during the data collection, we also averaged the different repeated measurements of the same antler. If only one photograph of an individual was obtained and this photograph contained the length of the antler on one side only, we used this length instead of the average value.

## 3. Ethical note

All research procedures were approved by the UCD Animal Research Ethics Committee (permit AREC-E-18-28). Capture and handling procedures prioritised animal welfare. Handlers consistently wore gloves to reduce the risk of human scent transfer, which could cause maternal rejection. To minimise stress, all handling was conducted in silence, and researchers promptly left the bed-site after each neonate’s release. An experienced wildlife biologist supervised the capture, handling, tagging, and sampling processes. Long-term monitoring of the tagging protocol has shown no negative impact on neonate survival in this population (Hayden, Moore, & Kelly, 1992).

## 4. Statistical Analysis

We conducted all analyses in R version 4.2.2 (R Core Team, 2022) using Bayesian multivariate mixed-effects models in the MCMCglmm package version 2.34 (Hadfield, 2010) and a structural equation model (SEM) in the sem package version 3.1-16 (Fox, 2006) to study the long-term effect of neonatal responses on male yearling antler length. Our analyses followed three key steps:

i. assessing the repeatability and correlations among heart rate, first reaction and second reaction at release,
ii. evaluating the effects of neonatal responses on growth rate across sexes, and
iii. examining the long-term effects of neonatal responses and growth rate on male yearling antler length.

The three steps implied running MCMCglmm models with different response variables and predictors (described in full in the corresponding 3 following sections of the methods). In general, for all models, we scaled all continuous variables and log-transformed heart rate to improve model convergence. We included neonate identity (neonate ID) as a random intercept and excluded rows with missing values in any explanatory variable from the analyses. For ordinal response variables (reactions), we constrained the residual variance to 1 (Hadfield, 2010). We ran each MCMC chain for 1,050,001 iterations, with a thinning interval of 1,000 and a burn-in period of 50,000 iterations. We assessed model convergence using the Heidelberger–Welch stationarity test (Heidelberger & Welch, 1983), visual inspection of trace plots, and examination of autocorrelation among samples.

For quantities derived from the MCMCglmm models, we report posterior means and 95% highest posterior density credible intervals (CI). We considered an association supported when its 95% CI excluded zero. (Allen, Street, & Capellini, 2017). We extracted the posterior means of the random intercepts (BLUPs) to visualise the relationships between neonatal responses and growth rate (Houslay & Wilson, 2017).

### 4.1 Model 1: Repeatability and correlations between neonatal responses

We fitted a trivariate Bayesian mixed effect model with heart rate, first reaction and second reaction as response variables. Prior to fitting the Bayesian multivariate mixed-effects models, we determined the optimal fixed-effect structure for each response variable separately using frequentist mixed-effects models, which provided a computationally tractable approach to model selection. For the ordinal response variables (first reaction and second reaction), we fitted cumulative link mixed models (CLMMs) using the ordinal package (Christensen, 2019), with individual identity as a random intercept. For heart rate, we fitted a linear mixed-effects model using lme4 package (Bates et al., 2015), again with individual identity as a random intercept. In all cases, we specified a full model including all candidate predictors: sex, capture time (hour), number of captures (capture), prior behaviour, body weight, bed-site visibility, air temperature, handling time and year. We then performed model selection using the MuMIn package (Barton & Barton, 2015). Models were ranked by Akaike Information Criterion (AIC), and the fixed-effect structure of the best-supported model (lowest AIC) was carried forward into the MCMCglmm models for each response. We included air temperature, capture time (hour), prior behaviour, weight, sex and capture year as predictors for heart rate; air temperature, prior behaviour, weight, number of capture and capture year for first reaction; and capture time (hour), prior behaviour, weight and number of captures for second reaction. The final model structure for the model is given in Supplement Table S1. In MCMCglmm models, we computed the correlation coefficients at the among-individual level and repeatability estimates, along with their 95% credible intervals, following Houslay and Wilson (2017). Since the first reaction and second reaction were highly collinear (see Results), we only used first reaction in the subsequent analyses.

### 4.2 Model 2: Sex-specific effects of neonatal responses on growth rate analysis

To test if neonatal responses show different effects on growth rate across sexes, we ran two separate trivariate mixed effect models for males and females with heart rate, first reaction and growth rate as response variables. We derived the growth rate variable in a two-step process. First, for all neonates captured at least twice with valid weight and age records, we fitted a linear mixed-effects model using the lme4 package version 1.1-31 (Bates et al., 2015; Griffin et al., 2022), with capture weight as the response variable. This model included linear and quadratic terms for age (days) as fixed effects and individual ID as a random effect (random intercept and slope for age). Second, we extracted the individual-specific random slope for age, representing individual growth rate, and its associated standard error for each individual. This growth rate estimate was then incorporated as a response variable in the trivariate model, with its standard error propagated as a measurement error variance via the mev argument.

We then ran two separate trivariate mixed-effects models for males and females, with heart rate, first reaction, and growth rate as response variables. The predictors for heart rate and first reaction were identical to Model 1, and the within-individual variance for growth rate was fixed to zero as it had no within-individual variance (Houslay & Wilson, 2017). Although including sex as a fixed effect in a single unified model is generally preferable, this approach is not appropriate in our specific modelling framework. In MCMCglmm, a fully unified model would instead require sex-stratified random effect covariance matrices, which introduces severe convergence challenges given our model complexity, mixed response variable families (Gaussian and ordinal), and the incorporation of measurement error variances. Separate sex-specific models are therefore used in this context. To formally test whether the among-individual correlations differed between sexes, we extracted the full posterior distributions of each correlation from both models and computed their difference. We computed the correlation coefficients at the among-individual level, along with their 95% credible intervals, following (Houslay & Wilson, 2017). We then computed the posterior distribution of the difference between the male and female correlations, and report its posterior mean and 95% CI. Consistent with our inference criterion used throughout, we considered the sex difference meaningful if less than 5% of the posterior distribution of the difference overlapped with zero.

### 4.3 Model 3: The long-term effect of neonatal responses on male yearling antler length analysis

To test if neonatal responses affect male yearling antler length directly or indirectly (via growth rate), we used a structural equation model (SEM). First, we ran a multivariate mixed-effects model (MCMCglmm) for males with heart rate, first reaction, growth rate, and antler length (at age one) as response variables. Predictors for heart rate and first reaction were identical to Model 1 (The final model structure for the model is given in Supplement Table S2). Within-individual correlations for growth rate and antler length were set to 0 since they had no within-individual variance (Houslay & Wilson, 2017). Second, we extracted the estimated among-individual correlation matrix from this model. This matrix was then used as the input for the SEM to evaluate the hypothesised paths between personality, growth rate, and antler length (Gaona-Gordillo et al., 2023).

## Results

### 1. Model 1: Repeatability and correlations between neonatal responses

A total of 708 individuals had complete records across all three response variables (heart rate, first reaction, and second reaction) and were included in the analysis across eight field seasons (2018–2025; 2018: n = 93; 2019: n = 80; 2020: n = 94; 2021: n = 94; 2022: n = 83; 2023: n = 83; 2024: n = 90; 2025: n = 91). Of these, 274 individuals had repeated measures (2018: n = 36; 2019: n = 45; 2020: n = 42; 2021: n = 27; 2022: n = 30; 2023: n = 25; 2024: n = 35; 2025: n = 34). All three neonatal response measures showed repeatable among-individual variation. Adjusted repeatability estimates were R = 0.208 (95% CI [0.143, 0.271]) for heart rate, R = 0.797 (95% CI [0.746, 0.850]) for the first reaction, and R = 0.519 (95% CI [0.450, 0.592]) for the second reaction. At the among-individual level, heart rate was positively correlated with both the first reaction (correlation coefficient (ri) = 0.381, 95% CI [0.200, 0.543]), and the second reaction (ri = 0.370, 95% CI [0.178, 0.527]). The first and second reaction were highly collinear (ri = 0.941, 95% CI [0.915, 0.965]). Based on this high collinearity, we used only the first reaction in subsequent analyses.

The model revealed that heart rate, first reaction, and second reaction are influenced by several factors: air temperature, prior behaviour, weight, year of capture, and sex affected heart rate; time of capture, prior behaviour, year of capture, and weight influenced the first reaction; prior behaviour and weight influenced the second reaction. Full model outputs are detailed in Supplementary Table S1.

### 2. Model 2: Sex-specific effects of neonatal responses on growth rate

Model 2 required complete records across two response variables only (heart rate and first reaction) rather than all three, meaning individuals with missing second reaction data but valid heart rate and first reaction records could be retained. We assessed sex-specific among-individual correlations between neonatal responses and growth rate using 154 males and 125 females, all captured on two or more occasions across eight field seasons (males: 2018: n = 22; 2019: n = 23; 2020: n = 27; 2021: n = 17; 2022: n = 17; 2023: n = 10; 2024: n = 20; 2025: n = 18; females: 2018: n = 14; 2019: n = 23; 2020: n = 16; 2021: n = 11; 2022: n = 13; 2023: n = 17; 2024: n = 16; 2025: n = 15). First reaction and growth rate showed a negative correlation at the among-individual level in males (r_i_ = -0.593, 95% CI [-0.796, -0.386]; Fig. 2a, 2c), but not in females (r_i_ = -0.257, 95% CI [-0.497, 0.055]; Fig. 2b, 2d). The male among-individual correlation was more negative than the female correlation (posterior probability = 0.969; pMCMC = 0.031), supporting a stronger negative association in males. In contrast, heart rate and growth rate were not correlated in either males (r_i_ = - 0.262, 95% CI [-0.510, 0.017]; Fig. 2a, 2e) or females (r_i_ = -0.198, 95% CI [-0.409, 0.068]; Fig. 2b, 2f). The correlations did not differ meaningfully between sexes (posterior probability = 0.624; pMCMC = 0.376), suggesting that heart rate showed no sex-specific relationship with growth rate. Table 1 presents the posterior means and 95% credible intervals for the explanatory variables used to estimate among-individual correlations between neonatal responses at capture and growth rate for both sexes.

**Figure. 2.**
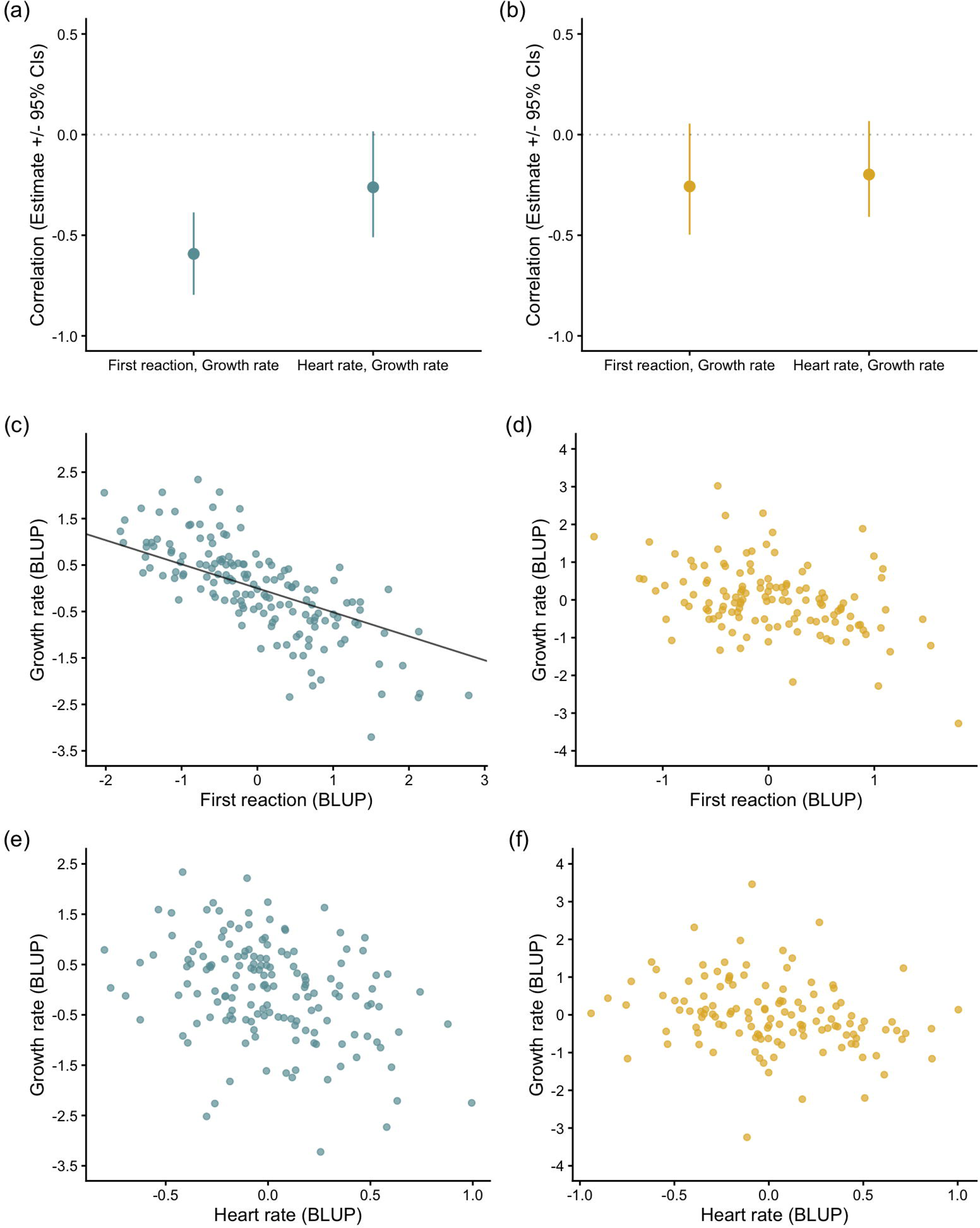
Among-individual correlations between heart rate at capture, first reaction at capture and growth rate in males (a) and females (b) from the multivariate model (MCMCglmm). Points represent posterior mean estimates of correlations, and vertical lines indicate 95% credible intervals (CIs). Correlations whose credible intervals do not overlap zero indicate supported associations. The relationships between first reaction and growth rate in males (c) and in females (d) and the relationships between heart rate and growth rate in males (e) and in females (f). Posterior means of the random intercepts (BLUPs) were used here for visualisation purposes only. Solid trendline indicates a supported effect.

**Table. 1.**
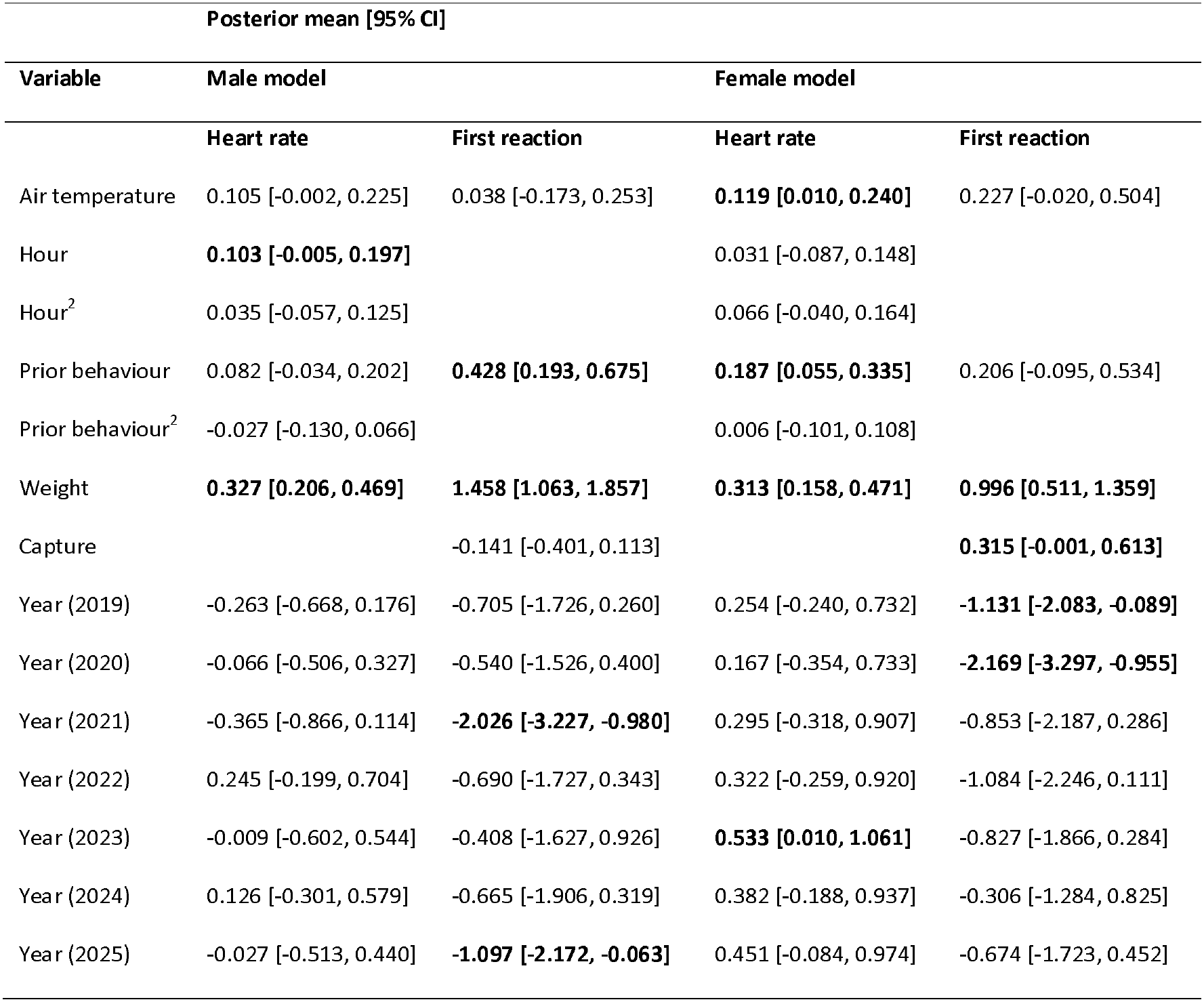
Structure and output of the trivariate model (MCMCglmm) used for the analysis of the among-individual correlations between heart rate at capture, first reaction at capture and growth rate in males and females. Posterior means with their associated 95% Credible Intervals of each of the explanatory variables included are given. Empty cells indicate that the explanatory variable was not included in the model for the respective response variables. Cells displayed in bold indicate supported associations.

|  | Posterior mean [95% CI] |  |  |  |
|---|---|---|---|---|
| Variable | Male model |  | Female model |  |
|  | Heart rate | First reaction | Heart rate | First reaction |
| Air temperature | 0.105 [-0.002, 0.225] | 0.038 [-0.173, 0.253] | **0.119 [0.010, 0.240]** | 0.227 [-0.020, 0.504] |
| Hour | **0.103 [-0.005, 0.197]** |  | 0.031 [-0.087, 0.148] |  |
| Hour<sup>2</sup> | 0.035 [-0.057, 0.125] |  | 0.066 [-0.040, 0.164] |  |
| Prior behaviour | 0.082 [-0.034, 0.202] | **0.428 [0.193, 0.675]** | **0.187 [0.055, 0.335]** | 0.206 [-0.095, 0.534] |
| Prior behaviour<sup>2</sup> | -0.027 [-0.130, 0.066] |  | 0.006 [-0.101, 0.108] |  |
| Weight | **0.327 [0.206, 0.469]** | **1.458 [1.063, 1.857]** | **0.313 [0.158, 0.471]** | **0.996 [0.511, 1.359]** |
| Capture |  | -0.141 [-0.401, 0.113] |  | **0.315 [-0.001, 0.613]** |
| Year (2019) | -0.263 [-0.668, 0.176] | -0.705 [-1.726, 0.260] | 0.254 [-0.240, 0.732] | **-1.131 [-2.083, -0.089]** |
| Year (2020) | -0.066 [-0.506, 0.327] | -0.540 [-1.526, 0.400] | 0.167 [-0.354, 0.733] | **-2.169 [-3.297, -0.955]** |
| Year (2021) | -0.365 [-0.866, 0.114] | **-2.026 [-3.227, -0.980]** | 0.295 [-0.318, 0.907] | -0.853 [-2.187, 0.286] |
| Year (2022) | 0.245 [-0.199, 0.704] | -0.690 [-1.727, 0.343] | 0.322 [-0.259, 0.920] | -1.084 [-2.246, 0.111] |
| Year (2023) | -0.009 [-0.602, 0.544] | -0.408 [-1.627, 0.926] | **0.533 [0.010, 1.061]** | -0.827 [-1.866, 0.284] |
| Year (2024) | 0.126 [-0.301, 0.579] | -0.665 [-1.906, 0.319] | 0.382 [-0.188, 0.937] | -0.306 [-1.284, 0.825] |
| Year (2025) | -0.027 [-0.513, 0.440] | **-1.097 [-2.172, -0.063]** | 0.451 [-0.084, 0.974] | -0.674 [-1.723, 0.452] |

### 3. Model 3: The long-term effect of neonatal responses on male yearling antler length

We used SEM to test for direct and indirect effects of neonatal responses on male yearling antler length in 73 males captured across seven field seasons (2018: n = 14; 2019: n = 18; 2020: n = 13; 2021: n = 12; 2022: n = 3; 2023: n = 5; 2024: n = 8). Note that no antler data were available for the 2025 cohort, as these individuals had not yet reached one year of age at the time of analysis. After incorporating the correlation matrix obtained from the multivariate mixed effects models (See supplement Table S2 for the full results), the SEM model supported a significant indirect pathway: neonatal behavioural reaction negatively affected growth rate, which in turn positively affected antler length (Figure 3, Table S3). These results imply that neonate with lower behavioural reaction scores grew faster and had longer yearling antler length. We found no evidence for a direct effect of behavioural reaction score or heart rate on antler length. Full model paths and estimates are provided in Supplementary Table S3.

**Figure. 3.**
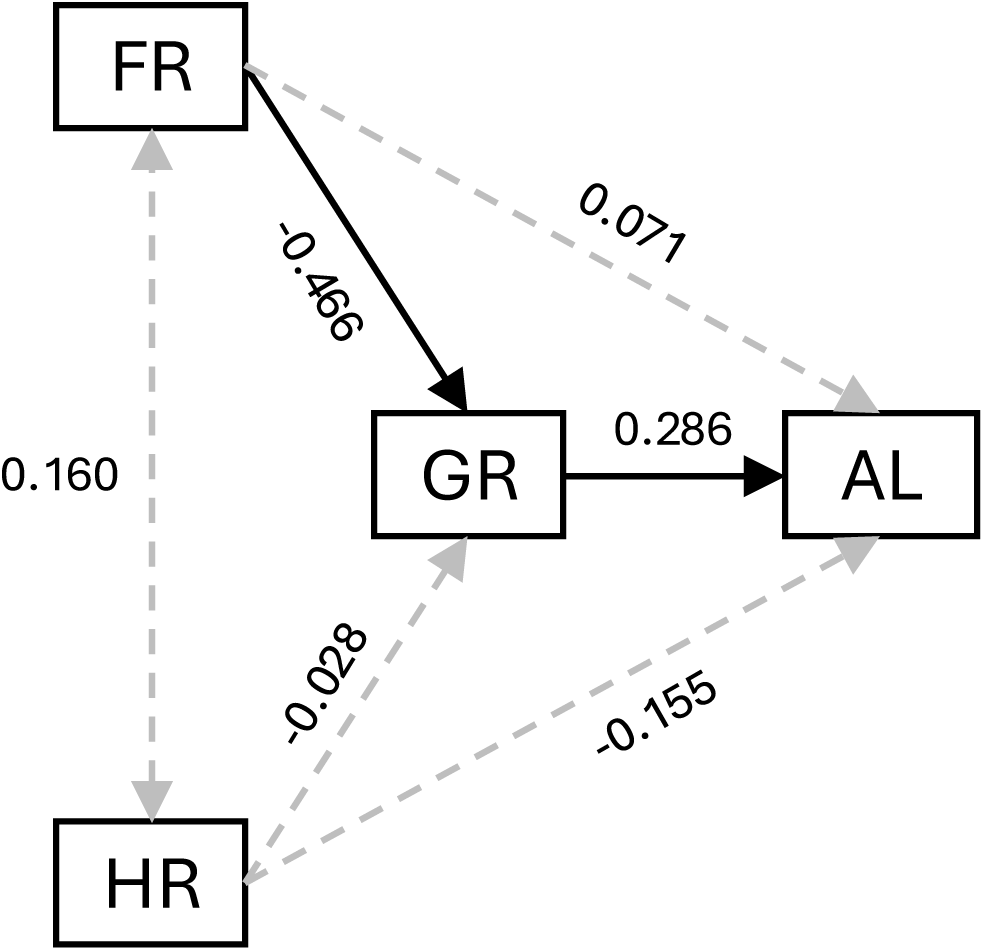
Visualisation of the results from structural equation model (SEM) used for estimating hypothesised paths between neonatal responses, growth rate and male antler length at age one. Shown are the point estimates for each of six structural paths connecting heart rate (HR), first behavioural reaction (FR), growth rate (GR) and antler length (AL) among individuals. Numbers correspond to standardised path coefficients (single-headed arrows) or residual among-individual correlations (double-headed arrows). Black and solid versus grey and dashed arrows are statistically significant (p < 0.05) versus non-significant (p > 0.05) estimates, respectively. See Table S3 for the uncertainty (SE) and statistical significance of each path.

## Discussion

This study investigated the effects of neonatal responses on developmental trajectories in a large mammal, fallow deer, and showed that stable among-individual differences in behaviour can be detected during the neonatal period and that their associations with development differed between females and males. Our results demonstrate three key findings: (1) Higher behavioural reaction scores were associated with slower early-life growth in males, whereas the corresponding association was not clearly supported in females. (2) Heart rate was not correlated with growth in either sex, suggesting that these two neonatal response measures differed in their association with growth at this life stage, and they are not interchangeable indicators of early-life developmental trajectories. (3) In the male structural equation model, behavioural reaction was negatively associated with yearling antler length indirectly through growth. These findings concern how behavioural, physiological, and morphological variation is organised during development, consistent with the component-of-variation perspective advocated by Réale, Dall, and Dingemanse (2026).

The negative among-individual correlation between behavioural reaction and growth indicates that males that were consistently more reactive during neonatal handling also tended to grow more slowly during early life. Comparable links between consistent behaviour and individual growth rate have been observed in other taxa, although their direction and expression vary among systems (Biro, Adriaenssens, & Sampson, 2014). Such associations between consistent behaviour and growth can arise when stable differences in individual state, including intrinsic growth capacity, influence behavioural expression, and may themselves be established early through genetic or persistent environmental effects (Biro, Adriaenssens, & Sampson, 2014). The stronger association in males may reflect the greater importance of rapid somatic growth during male development. In sexually dimorphic ungulates, males follow a more growth-intensive developmental trajectory as differences in body size and male-specific morphology emerge (Pérez-Barbería, Gordon, & Pagel, 2002). Consequently, early differences among males may become more clearly expressed in their growth rates, strengthening the covariance between neonatal behaviour and subsequent development. The sex difference therefore suggests that the organisation of behavioural and developmental variation is already sex-specific during early life.

Our study also indicates that heart rate and behavioural reaction captured related but non-equivalent aspects of neonatal response. It suggested a more complex connection between personalities and growth, which aligns with existing mixed findings on the relationship between an individual’s behavioural response to risk (coping style) and its physiological stress response to risk (Clary et al., 2014; Eleftheriou, Williams, & Luis, 2022), with evidence showing that they should be treated differently in some species (Lapointe et al., 2015; Qu et al., 2018). Previous research on this fallow deer population found that a slower heart rate in neonates is linked to higher survival (Amin, Verbeek, et al., 2022), while behavioural response to human handling is negatively linked to trappability and vigilance behaviour (Amin, Jennings, et al., 2022; Yu et al., 2025). These findings imply that these two aspects of personality should be considered separately and suggest that different personality components may be linked to different life-history traits (Smith & Blumstein, 2008). More generally, this distinction illustrates how the variance and covariance associated with different traits can inform our understanding of phenotypic organisation during development (Réale, Dall, & Dingemanse, 2026).

The male structural equation model extended the association between neonatal behaviour and early growth to yearling morphology. Behavioural reaction was not directly associated with antler length, but males with higher reaction scores tended to grow more slowly, and faster-growing males had longer antlers at one year. This pattern likely arose because growth provides the developmental connection between early behavioural variation and later morphology. Behavioural reaction and growth may be components of an integrated phenotype established early in life, such that individuals differ consistently in both their responses to handling and their subsequent growth trajectories. In males, these growth trajectories are expressed in antler development because antlers are rapidly growing and are closely linked to somatic development (Ciuti & Apollonio, 2011; Vanpé et al., 2007). This association may therefore be especially apparent in a strongly sexually dimorphic species such as fallow deer, in which male growth contributes to the development of traits that later influence social status and mating success (Ciuti & Apollonio, 2011). The indirect association therefore places neonatal behavioural variation within a broader pattern of male phenotypic development, linking early behavioural organisation with growth and the emergence of sexually dimorphic morphology.

In conclusion, this study shows that repeatable among-individual differences in behaviour and heart rate were already detectable during the neonatal period in wild fallow deer. Their developmental associations were not uniform. Behavioural reaction was associated with early growth in males, evidence for the corresponding female association was weaker, and heart rate was not associated with growth in either sex. In males, the association between neonatal behavioural reaction and yearling antler length was expressed indirectly through growth within the hypothesised model. Together, these results show that early behavioural variation is associated with the organisation of developing phenotypes, with stronger evidence for this developmental association in males. Following individuals further into adulthood will help determine how long these neonatal differences persist and whether the observed developmental associations extend to later morphology and performance.

