## supplementary methods for "Neonatal personality is associated with sex-specific developmental trajectories in a wild mammal"

***The capture protocol***

Following long-term herd management protocols established in the early 1970s, fawns were captured and tagged with unique ear tags (Hayden, Moore, & Kelly, 1992). Traditional fawning grounds were divided into seven sectors; these were patrolled by a team of approximately 15 trained personnel under the supervision of an experienced biologist. Sectors were monitored on a rotational basis, ensuring each was visited at least once every three days. Fawns were secured using long-handled circular fishing nets. To evaluate the repeatability of behavioural measures, opportunistic recaptures were conducted, avoiding same-day recaptures.

Upon capture, fawn age was estimated based on the condition of the umbilical cord, the shoe and dew claws (Galli et al., 2008), and body mass was recorded using a 100-liter bag and digital scale. We assessed bed-site visibility (sensu (Bongi et al., 2008)) by placing a fawn-sized cardboard target (45 × 36 cm) at the capture location. Visibility was calculated as the proportion of the target visible from 10 m away in the four cardinal directions at an observer height of 70 cm, which simulates the perspective of a red fox (*Vulpes vulpes*). Air temperature at the bed-site was also recorded.

Bongi, P., Ciuti, S., Grignolio, S., Del Frate, M., Simi, S., Gandelli, D., & Apollonio, M. (2008). Anti‐predator behaviour, space use and habitat selection in female roe deer during the fawning season in a wolf area. *Journal of Zoology*, *276*(3), 242-251.

Galli, A., Focardi, S., Iseppi, M., & Piasentier, E. (2008). Age determination in fallow deer Dama dama neonates. *Wildlife Biology*, *14*(3), 386-390.

Hayden, T., Moore, N., & Kelly, P. (1992). The fallow deer of Phoenix Park: an evolving management plan. Proc. 2nd Deer Park Symp.: management, welfare and conservation of park deer,
