## Supplement Table for "Neonatal personality is associated with sex-specific developmental trajectories in a wild mammal"

Table. S1 Structure and output of the multivariate model (MCMCglmm) used for the analysis of the repeatability and among-individual correlations between heart rate at capture, first reaction at capture and second reaction at capture in fallow deer neonates. Posterior means with their associated 95% Credible Intervals of each of the explanatory variables included are given. Empty cells indicate that the explanatory variable was not included in the model for the respective response variables.

|  | **Posterior mean [95% CI]** |  |  |
| --- | --- | --- | --- |
| **Variable** | **Heart rate** | **First reaction** | **Second reaction** |
| Air temperature | 0.127 [0.072, 0.181] | 0.182 [0.041, 0.332] |  |
| Hour | 0.066 [0.000, 0.124] |  | 0.080 [-0.026, 0.179] |
| Hour^2^ | 0.031 [-0.020, 0.081] |  |  |
| Prior behaviour | 0.251 [0.182, 0.325] | 0.472 [0.287, 0.642] | 0.321 [0.182, 0.444] |
| Prior behaviour^2^ | -0.011 [-0.060, 0.038] |  |  |
| Weight | 0.213 [0.143, 0.298] | 1.397 [1.159, 1.678] | 0.777 [0.607, 0.925] |
| Capture |  | 0.081 [-0.069, 0.246] | 0.007 [-0.088, 0.119] |
| Sex (m) | 0.275 [0.154, 0.376] |  |  |
| Year (2019) | 0.129 [-0.096, 0.350] | -0.756 [-1.437, 0.050] |  |
| Year (2020) | 0.185 [-0.044, 0.409] | -0.863 [-1.558, -0.206] |  |
| Year (2021) | 0.148 [-0.089, 0.367] | -0.875 [-1.546, -0.196] |  |
| Year (2022) | 0.450 [0.250, 0.690] | -1.019 [-1.695, -0.284] |  |
| Year (2023) | 0.320 [0.108, 0.573] | -0.745 [-1.447, 0.059] |  |
| Year (2024) | 0.387 [0.181, 0.613] | -0.195 [-0.941, 0.425] |  |
| Year (2025) | 0.423 [0.209, 0.657] | -0.334 [-1.111, 0.316] |  |

Table. S2 Structure and output of the multivariate model (MCMCglmm) used for the analysis of the among-individual correlations between heart rate at capture, first reaction at capture, growth rate and antler length at age one in male fallow deer. Posterior means with their associated 95% Credible Intervals of each of the explanatory variables included are given. Empty cells indicate that the explanatory variable was not included in the model for the respective response variables.

|  | **Posterior mean [95% CI]** |  |
| --- | --- | --- |
| **Variable** | **Heart rate** | **First reaction** |
| Air temperature | 0.122 [-0.043, 0.272] | 0.001 [-0.278, 0.309] |
| Hour | 0.084 [-0.057, 0.225] |  |
| Hour^2^ | 0.058 [-0.061, 0.201] |  |
| Prior behaviour | 0.019 [-0.124, 0.189] | 0.495 [0.180, 0.820] |
| Prior behaviour^2^ | 0.018 [-0.113, 0.159] |  |
| Weight | 0.460 [0.293, 0.636] | 1.239 [0.774, 1.777] |
| Capture |  | -0.153 [-0.504, 0.275] |
| Year (2019) | -0.508 [-1.068, 0.044] | -0.459 [-1.696, 0.566] |
| Year (2020) | -0.159 [-0.877, 0.473] | -0.844 [-2.270, 0.473] |
| Year (2021) | -0.380 [-1.025, 0.291] | -1.881 [-3.419, -0.541] |
| Year (2022) | 0.594 [-0.446, 1.565] | -0.329 [-2.369, 1.842] |
| Year (2023) | -0.246 [-1.010, 0.543] | -0.676 [-2.229, 1.075] |
| Year (2024) | 0.268 [-0.408, 1.051] | -0.552 [-2.002, 0.851] |

Table. S3 Output of the structural equation model used for estimating hypothesised paths between heart rate at capture, first reaction at capture, growth rate and antler length at age one in male fallow deer. The model was fitted using the among-individual correlation matrix among all four mean- and variance-standardised traits from MCMCglmm presented in Table S2. We printed each path’s standardised estimate, standard error, z value, and p-value. We present two types of parameters: standardised path coefficients (→) and correlations (↔).

| **Path** | **Estimate** | **Standard error** | **z value** | **p value** |
| --- | --- | --- | --- | --- |
| First reaction → Growth rate | -0.466 | 0.105 | -4.424 | <0.001 |
| Heart rate → Growth rate | -0.028 | 0.105 | -0.267 | 0.789 |
| First reaction → Antler length | 0.071 | 0.128 | 0.557 | 0.578 |
| Heart rate → Antler length | -0.155 | 0.113 | -1.364 | 0.172 |
| Growth rate → Antler length | 0.286 | 0.127 | 2.254 | 0.024 |
| First reaction ↔ Heart rate | 0.160 | 0.119 | 1.340 | 0.180 |
